# Recent social loss, not chronic isolation, reshapes sleep in *Drosophila*

**DOI:** 10.64898/2026.07.12.738068

**Authors:** Binbin Wu, William W. Ja

**Affiliations:** Department of Neuroscience, The Herbert Wertheim UF Scripps Institute for Biomedical Innovation & Technology, Jupiter, FL 33458, USA

## Abstract

Social isolation is associated with altered sleep. Although *Drosophila melanogaster* is widely used to study social experience, it remains unclear whether sleep changes attributed to isolation reflect chronic deprivation or the consequences of a recent loss of social contact. Conventional isolation paradigms alter social context at the time of testing, making these possibilities difficult to distinguish. Here, we developed sociSleep, an identity-preserving tracking system that quantifies sleep without inducing social-state transitions. Using this approach, we found that chronic social isolation had little effect on sleep, whereas recent social loss robustly increased sleep. Importantly, this effect is not attributable to sleep loss or baseline differences between solo and group housing: in the absence of a social-state transition, solo- and socially housed flies exhibit equivalent sleep. These findings show that recent social experience, rather than chronic isolation alone, is a major modulator of sleep in flies. This distinction provides new insight into the relationship between social connectedness, loneliness and sleep health.

## Introduction

*Drosophila melanogaster* has emerged as a powerful model for dissecting the genetic and neural mechanisms underlying social behavior, including courtship, aggression, and pheromone-mediated communication^1–3^. Social experience can modify many these behaviors^4^, raising a broader question of whether it also influences fundamental behavioral states. Sleep is particularly well suited for addressing this question because it is an evolutionarily conserved behavior and can be quantified in *Drosophila* with established criteria^5^. In humans, social isolation and loneliness are associated with adverse health outcomes and behavioral changes^6,7^, including disrupted sleep^8,9^. Whether social experience similarly shapes sleep in *Drosophila* remains unclear.

Previous studies in flies have reported that chronic isolation reduces sleep^10^ and that social enrichment increases it^11,12^, suggesting that sleep is sensitive to social experience. However, these paradigms typically change social context again at the time of measurement, by transferring group-housed flies into isolation for behavioral testing while maintaining chronically isolated flies in their original housing condition (Fig. 1A). As a result, behavioral differences may reflect not only prior social history, but also the immediate consequences of a recent transition in social state. More broadly, the assumption that behavioral phenotypes remain stable across such assay-imposed transitions has not been directly tested^10,11,13,14^.

**Figure 1:**
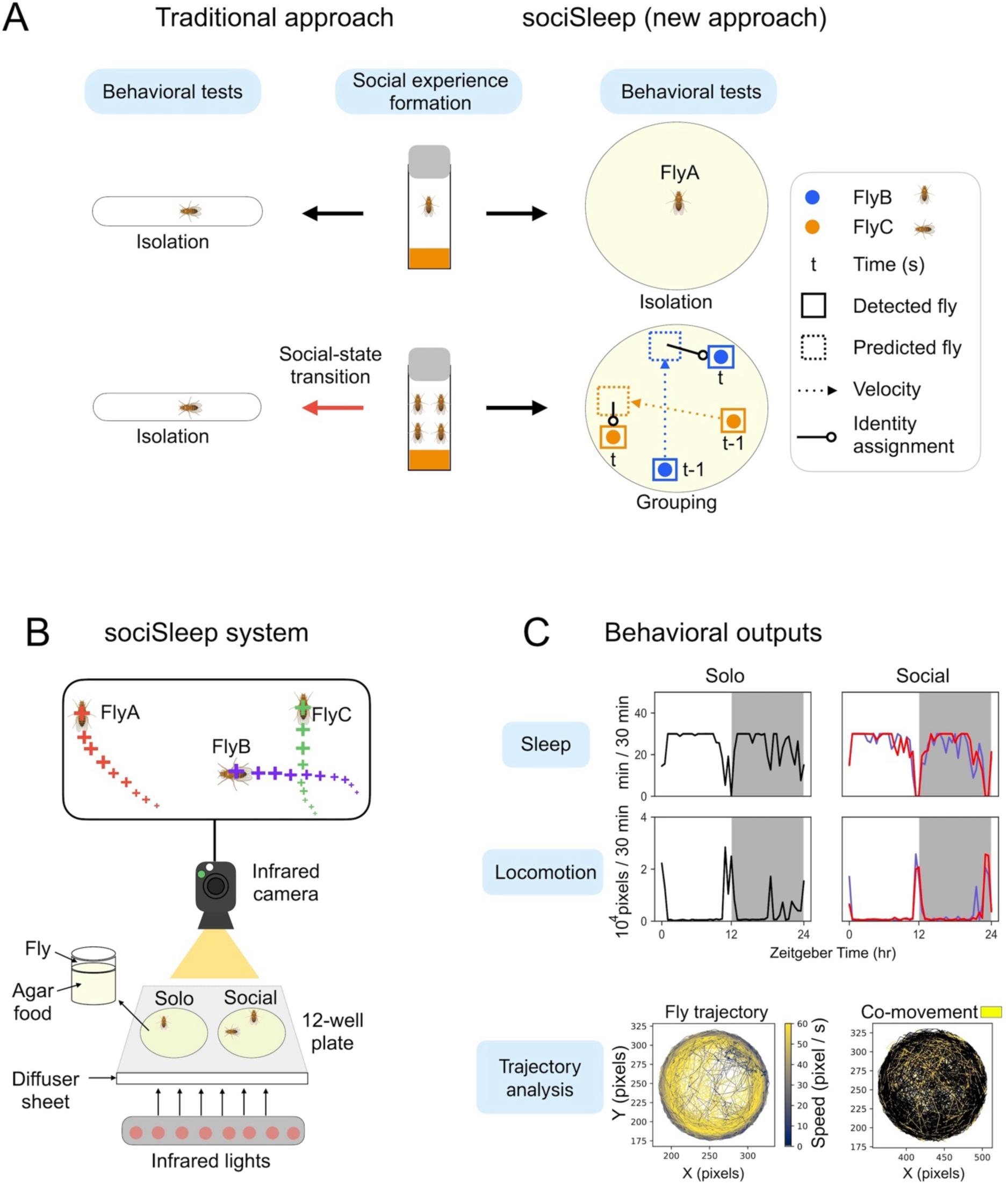
sociSleep, a new approach for studying social isolation impacts. (A) The commonly used experimental design (left) requires transferring flies from group housing to isolation for behavioral measurements, overlooking the effects of acute changes in social state. The new system (right) described here, sociSleep, enables continuous quantification of fly behavior without perturbing social state, minimizing confounding effects arising from behavioral responses to abrupt social-state transitions. This system assigns fly identities (FlyB and FlyC) at each time point (t) based on predicted positions derived from their coordinates and velocities at the previous time step (t-1). (B) Schematic of the sociSleep hardware assembly. The two central wells of a 12-well suspension culture plate serve as fly arenas and are filled to ∼ 90% volume with optimized agar-based food to provide nutrition, maintain humidity, and minimize reflections. A diffuser sheet is used to soften infrared illumination for reducing signal noise. Windows PC and Microsoft infrared cameras are recommended to ensure compatibility and facilitate camera parameter adjustment. (C) Representative behavioral outputs quantified by sociSleep for flies housed in either solo or social arenas. Top panels show sleep and locomotion profiles across 24 hr, expressed as sleep (min) or locomotor distance (10^4^ pixels) per 30-min bin. In the group arena, the blue and red lines represent the sleep or locomotor activity of two individual flies tracked simultaneously within the same arena. Bottom left, trajectory analysis of a single fly in a solo arena, with trajectory segments color-coded according to instantaneous movement speed. Bottom right, trajectories of two flies tracked in a social arena, with periods of co-movement highlighted in yellow. Co-movement is defined as epochs during which both flies are not immobile and remain within 30 pixels of each other.

Resolving this issue requires measuring behavior without altering the social conditions under which it is expressed, but that has been limited by the lack of reliable, accessible methods for long-term, identity-preserving tracking of individual flies within social groups. To address this problem, we developed sociSleep, a tracking system that enables continuous quantification of individual sleep and locomotor behavior in stable social environments over multiple days. This approach allows us to distinguish the effects of chronic isolation from those of recent social loss while preserving the social context in which behavior is expressed.

## Results

### Development of sociSleep

To quantify sleep in socially housed flies without changing their social context, we developed sociSleep, an identity-preserving tracking system for long-term behavioral monitoring in shared arenas. The system combines Kalman filter-based state prediction^15^ with nearest-neighbor assignment to maintain individual identities (Fig. 1A) and was designed for straightforward deployment with standard laboratory equipment (Fig. 1B). In addition to sleep, sociSleep extracts multiple behavioral metrics from tracked trajectories, including locomotor activity, movement speed, and close-proximity interactions (Fig. 1C).

Manual validation of randomly selected tracking sessions showed high agreement with automated scoring, with occasional identity swaps and brief signal-loss events (Table S1; Movies S3-S9). Because these events occurred during brief close-contact interactions when flies were active, they did not affect sleep detection, which was defined by sustained immobility, and statistical analyses were performed on chamber-level mean values. Together, these results establish sociSleep as a practical platform for quantifying sleep and other behavioral metrics in stable social environments without introducing the state transitions that can confound conventional isolation paradigms.

### *Drosophila* sleep shows limited sensitivity to chronic social isolation

Using sociSleep, we asked whether chronic social isolation alters sleep when social state remains constant during the measurement. To compare chronic isolation with continued social presence, white-eyed Canton-S (*w*CS) males were maintained under single- or group-housing conditions for varying durations, then transferred into sociSleep arenas either individually or in pairs, and tracked continuously across three 12:12-hr light:dark (LD) cycles (Fig. 2A). Although the social arenas contained fewer flies than the original group-housed vials, the smaller arena volume partially offset the reduced group size. Using a simple density-normalized encounter index, estimated pairwise interaction opportunities were comparable between group-housing vials and sociSleep social arenas (see Methods).

**Figure 2:**
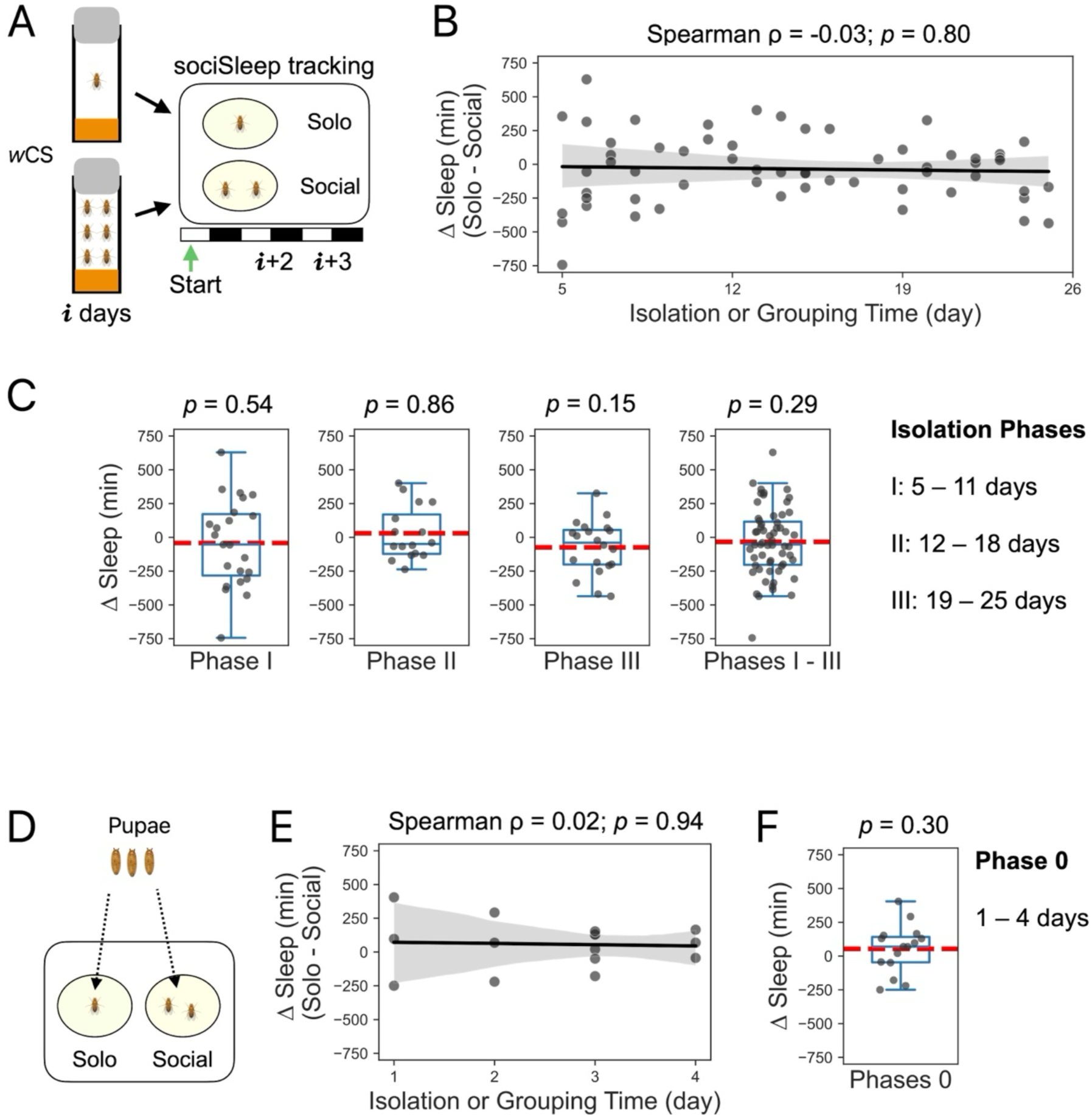
***Drosophila* sleep shows limited sensitivity to chronic or stable social contexts.** (A) Experimental design for examining the effects of chronic social isolation on sleep. ΔSleep was calculated for each trial as the difference in sleep amount between flies monitored in social arenas (two flies per arena) and solo arenas (one fly per arena). Flies were acclimated to the sociSleep system starting on day **i**+1, and behavioral data from the following two days (**i**+2 and **i**+3) were analyzed. (B) Correlation between ΔSleep and the duration of social isolation. No significant association was detected using the Spearman’s rank correlation. The solid line was made using robust linear regression, revealing that the slope is not significantly different from zero (*p* = 0.75). N = 30 independent paired trials. Each trial yielded one ΔSleep value per day through (solo sleep – averaged social sleep). (C) Statistical analysis of ΔSleep across different isolation phases. *P* values were calculated using the two-tailed Wilcoxon signed-rank test to determine if ΔSleep was different from zero. Data are presented as boxplots, in which the box represents the interquartile range (middle 50% of the data) and whiskers extend to 1.5 times of the interquartile range. The red line indicates the mean. (D) Experimental design for evaluating potential early effects of social isolation. Newly eclosed flies were immediately assigned to isolated or group-housed conditions. (E) Correlation between ΔSleep and the duration of social isolation. No significant association was detected using the Spearman’s rank correlation. The solid line was made using robust linear regression, revealing that the slope is not significantly different from zero (*p* = 0.87). N = 9 independent paired trials. (F) Sleep differences during the early isolation period. No significant difference in sleep was detected using the two-tailed Wilcoxon signed-rank test. Data are presented as boxplots, in which the box represents the interquartile range (middle 50% of the data) and whiskers extend to 1.5 times of the interquartile range. The red line indicates the mean.

Across 5–25 days of isolated and group housing, sleep differences showed no significant correlation with isolation duration (Spearman ρ = -0.03, *p* = 0.80), indicating that chronic exposure to distinct social conditions did not lead to progressive sleep divergence (Fig. 2B). Significant sleep differences were also not detected when the data were parsed into different phases from I (5–11 days) to III (19–25 days), and the combined data did not differ from zero (Fig. 2C). To test whether any effects can emerge earlier, in the first few days of adulthood, newly eclosed flies were assigned immediately to isolated or pair-housed conditions to minimize prior social experience (Fig. 2D). Again, no significant correlation (Spearman ρ = 0.02, *p* = 0.94) or group difference was detected (Fig. 2E-F).

Together, these data indicate that chronic isolation has limited effects on sleep when flies are measured without a recent change in social state, raising the possibility that previously reported effects arise from recent social-state transitions.

### Recent social loss increases sleep

Because traditional isolation paradigms change social context at the time of testing, we next asked whether recent social loss itself, rather than chronic isolation, is sufficient to alter sleep. We mimicked the housing change used in those paradigms by transferring group-housed *w*CS males to individually housed arenas, while control flies from the same vials were transferred in pairs to preserve social interactions after transfer (Fig. 3A). Flies that experienced recent social loss showed increased sleep on LD2 relative to socially housed controls, and the effect was attenuated on LD3 (Fig. 3B, D). This increase was not accompanied by a change in locomotor activity (Fig. 3C, E). Instead, trajectory-based analysis showed that singly housed flies moved faster, consistent with increased sleep without reduced locomotor distance (Fig. 3F). These results indicate that recent social loss is sufficient to alter sleep behavior and likely contributes to effects previously attributed to chronic social isolation in *Drosophila*. Consistent with this interpretation, baseline sleep was similar when flies were measured alone or in pairs in the absence of recent social loss (Fig. 3G-I), demonstrating that increased sleep is linked to the transition itself rather than to the immediate social condition during measurement.

**Figure 3:**
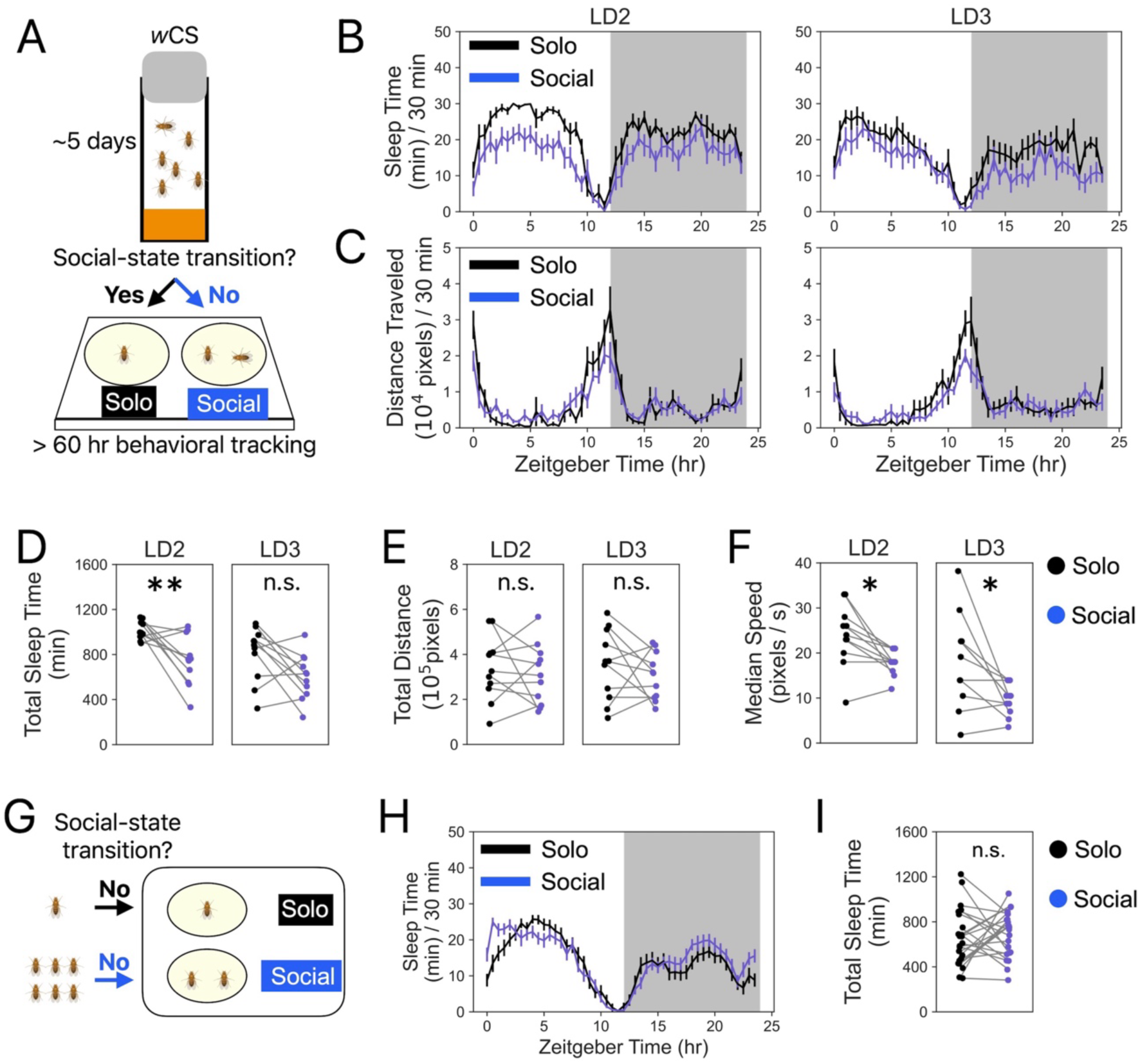
Recent social loss increases *Drosophila* sleep. (A) Schematic of the experimental design testing the effects of social-state transitions from group housing to isolation on sleep. Adult males were group-housed in vials for ∼5 days and then assigned to either solo (1 fly) or social (2 flies) arenas. Individuals were tracked using the sociSleep system for over 60 hr, and data from two consecutive days (LD2–LD3) were analyzed. (B) Sleep time (min) per 30 min across two days. For social arenas, values represent the mean of the two flies within each arena. (C) Locomotor distance (pixels) per 30 min across two days. Each fly yielded a distance value per day. For social arenas, values represent the mean of the two flies within each arena. (D) Daily sleep time in solo versus social arenas. (E) Daily locomotor distance in solo versus social arenas. (F) Daily median speed in solo versus social arenas. N = 11 independent paired trials. (G-I) Reanalysis of the Phase I cohort shown in Figure 2C using the raw sleep data. (G) Flies that remained isolated throughout the experiment were recorded individually (Solo), whereas flies that remained group-housed were recorded in pairs (Social), allowing comparison under stable social conditions without introducing a new transition at measurement. (H) Sleep time (min) per 30 min across the day. (I) Total daily sleep time (min). Data shown as mean ± s.e.m. Each pair of dots is connected by a gray line to represent one independent paired trial. Wilcoxon signed-rank tests were used for examining any significant differences between these paired measurements (two-tailed): n.s., *p* > 0.05; *, *p* < 0.05; **, *p* < 0.01.

### Brief social experience reshapes subsequent sleep

The finding that recent social loss increases sleep raises the question of how much social experience is required for an animal to “feel” socially deprived. To address this, we asked whether a brief, minimal period of social housing is sufficient to alter subsequent sleep when flies are returned to isolation. Previously single-housed flies were maintained for 24 hr either individually, in pairs, or in groups of eight, after which individual sleep was measured using a modified ARC system^16^ (Fig. 4A). This design both fixed the duration of social exposure (24 hr) and minimized the amount of social interaction possible by testing the smallest feasible group size (two flies per vial).

**Figure 4.**
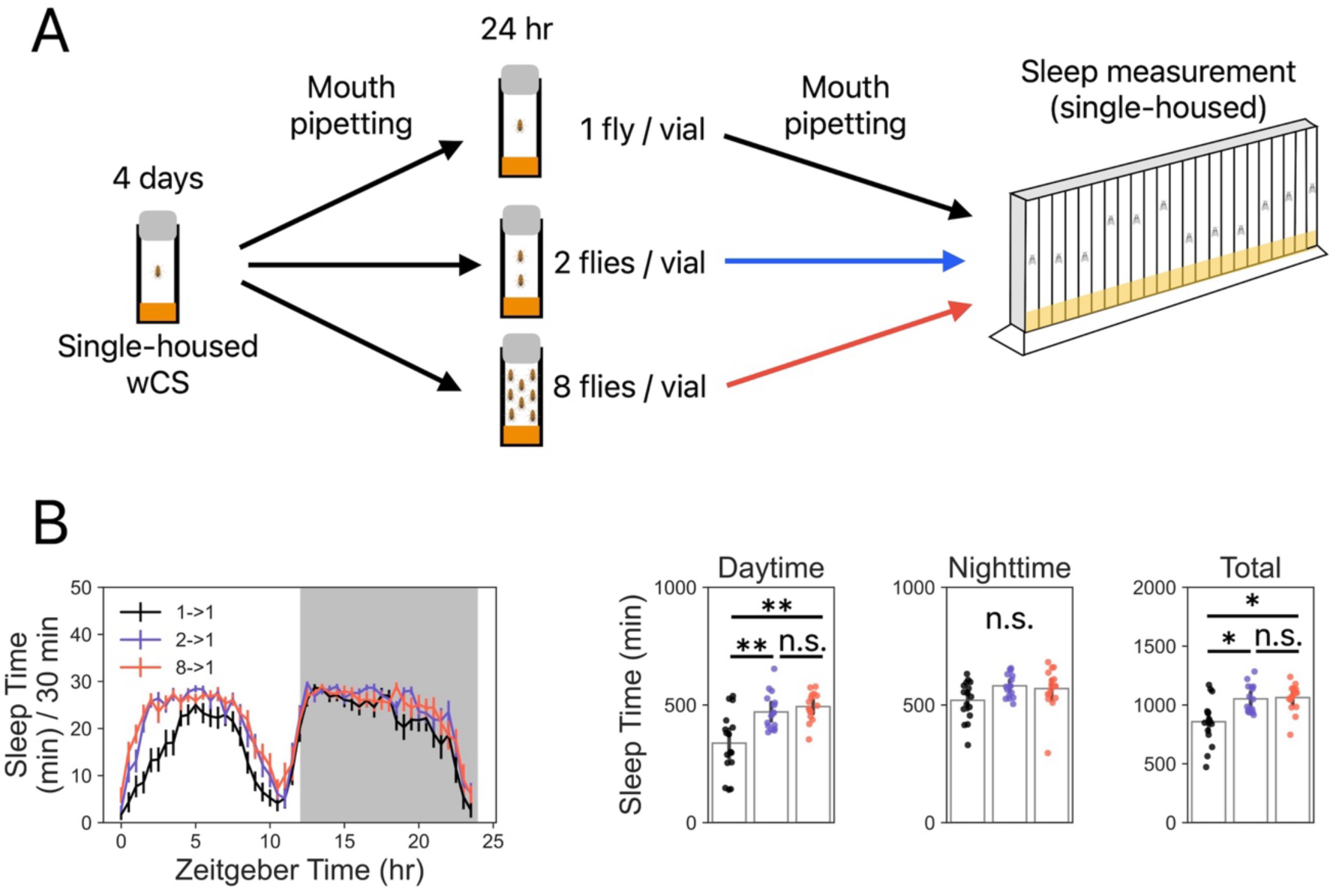
Brief social experience reshapes subsequent sleep. (A) Experimental design. Flies were individually housed for 4 days and then transferred to new vials for 24 hr either individually, in pairs, or in groups of eight. All flies were subsequently transferred to ARC chambers (1 fly/tube) for sleep measurement. (B) Sleep following 24 hr of varying social exposures. Flies that experienced 24 hr of social housing before sleep measurement (2◊1 and 8◊1) showed increased daytime and total sleep compared with flies maintained individually throughout (1→1). Sleep did not differ significantly between the 2◊1 and 8◊1 conditions. Data are shown as mean ± s.e.m. N = 16. Statistical comparisons were performed using two-tailed Wilcoxon signed-rank tests with Holm correction for multiple comparisons. n.s., *p* > 0.05; *, *p* < 0.05; **, *p* < 0.01.

Flies that experienced 24 hr of social housing showed increased sleep when tested in isolation compared with flies maintained individually throughout (Fig. 4B). Sleep increases were comparable for flies previously housed in pairs and in groups of eight, indicating that the effects does not scale simply with social group size over this interval. Together, these findings demonstrate that a brief, minimal period of social experience is sufficient to rapidly reshape subsequent sleep, without requiring prolonged or densely populated social housing.

### Social state-dependent sleep recovery

Although immediate social context had little effect on baseline sleep in the absence of recent social loss, it remained unclear whether flies would respond differently under elevated sleep pressure. Because traditional isolation paradigms eliminate ongoing social interactions during behavioral testing, we next asked whether these changes affect the expression of sleep rebound after sleep deprivation. Group-housed flies were either sleep deprived or left undisturbed, and recovery was measured in sociSleep under solo-housed or social-housed conditions. Flies recovering alone showed a robust sleep rebound during both day and night, whereas flies recovering in pairs showed only a modest daytime increase that did not reach significance (*p* = 0.098; Fig. 5A-B).

**Figure 5:**
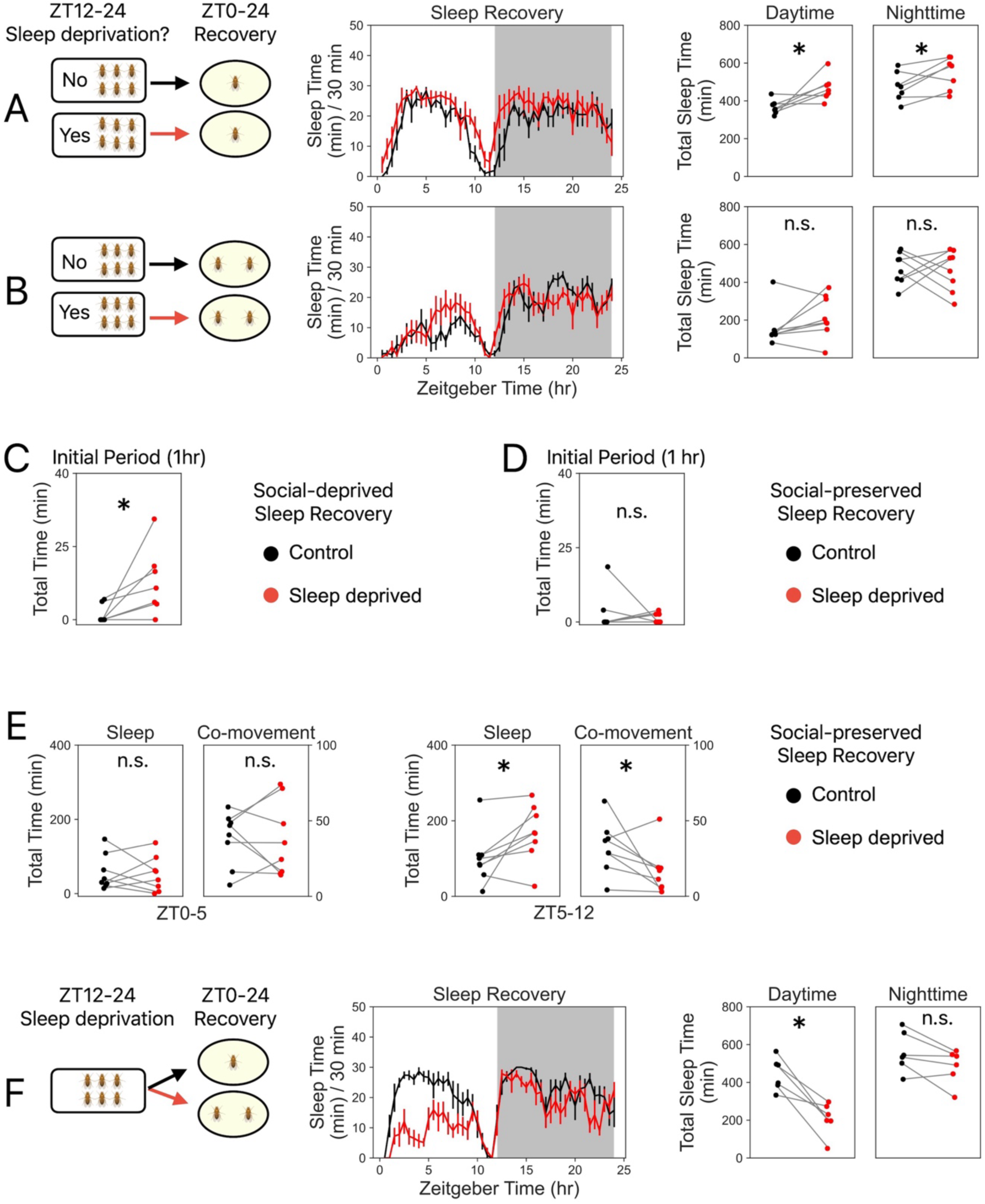
Current social state shapes responses to sleep loss. (A) Sleep recovery in individual flies after overnight (12 hr) sleep deprivation or no deprivation in group-housed vials, measured in solo-housed sociSleep arenas. N = 7 independent paired trials. (B) Sleep recovery after the same deprivation paradigm when recovery was measured in social-housed sociSleep arenas containing two flies. N = 8 independent paired trials. (C-D) Sleep during the first hour of recovery (ZT0-1) in social-deprived (C) and social-preserved (D) flies. (E) Sleep and co-movement (close-proximity interactions) during the early (ZT0-5) and late (ZT5-12) daytime recovery periods in social-housed sociSleep arenas. Co-movement was defined in Figure 1C and quantified as the cumulative duration (min) of all co-movement events. (F) Independent recovery experiment in which sleep-deprived flies from the same group-housing vials were assigned to recover either individually or in pairs. Isolated flies showed greater daytime sleep than paired flies. N = 6 independent paired trials. For 24-hr sleep curves, data are shown as mean ± s.e.m. For paired-dot plots, each pair of dots represents one independent paired trial and is connected by a gray line. Statistical significance in panels A-E was determined using one-tailed Wilcoxon signed-rank tests based on directional hypotheses regarding sleep rebound and co-movement. Panel F was analyzed using a two-tailed Wilcoxon signed-rank test for examining any significant differences between these paired total sleep measurements. n.s., *p* > 0.05; *, *p* < 0.05.

To determine whether this difference emerged immediately, we quantified sleep during the first hour of recovery. Social-deprived flies showed a rapid increase in sleep, whereas social-preserved flies did not (Fig. 5C-D). Instead, these social-preserved flies maintained close-proximity interactions during the early recovery period and increased sleep only later, as those interactions declined (Fig. 5E). These results suggest that ongoing social interaction delays sleep rebound after deprivation.

We then repeated the recovery experiment with a direct solo-versus-social comparison using only sleep-deprived group-housed flies. In this independent cohort, isolated flies again showed greater daytime sleep than paired flies (Fig. 5F). Thus, the sleep increase associated with recent social loss is observed not only under baseline conditions but also during recovery from a physiological challenge.

## Discussion

Social environment is often treated as a stable background variable, but our results show that recent changes in social state can themselves be a major determinant of behavior. By measuring sleep without perturbing social context, sociSleep allowed us to separate the effects of chronic isolation from those of social-state transition and reveal that the latter exerts the stronger influence on sleep phenotypes.

This distinction helps clarify how social experience shapes sleep in *Drosophila*. When social state was held constant during measurement, chronic isolation produced little evidence of progressive sleep disruption across days. By contrast, transfer from a social environment to isolation induced a rapid increase in sleep, and this effect was not explained simply by whether flies were measured alone or in pairs, because baseline sleep was similar across those conditions when recent social loss was absent (Fig. 3G-I). Together, these findings argue that recent social loss is a more powerful determinant of sleep than chronic isolation alone.

The broader implication is that behavioral phenotypes attributed to isolation may, in some cases, reflect the consequences of state transition rather than the accumulated effects of prolonged social deprivation. In flies, this issue is especially important because many commonly used paradigms alter social context at the time of assay. Our results therefore suggest that studies relying on post-housing transition may overestimate the effects of chronic isolation while underappreciating the impact of recent social loss.

This framework may extend beyond *Drosophila*. In mammals, acute isolation can rapidly alter subsequent social behavior, and recent work has identified neural circuits that contribute to the dynamic control of social homeostasis^17,18^. Although these studies did not address sleep directly, they support the broader idea that recent social loss can induce a distinct internal state with rapid behavioral consequences. The present results raise the possibility that some behavioral effects linked to isolation across species may depend not only on the duration of social deprivation but also on the timing of the most recent social transition.

Our sleep-deprivation experiments extend this conclusion by showing that immediate social context also shapes the expression of sleep homeostasis. Flies recovering alone displayed a robust sleep rebound, whereas flies recovering in pairs maintained close-proximity interactions during the initial recovery period. Sleep increased later in these socially recovering flies as those interactions declined, suggesting that ongoing social engagement can temporarily delay the behavioral expression of sleep pressure rather than eliminate sleep need altogether.

These findings also broaden the significance of the study beyond baseline sleep. Social state did not merely modulate a steady-state behavioral output; it influenced how flies responded to an internal homeostatic challenge. This result supports a view in which sleep expression is dynamically gated by current social conditions, allowing behavior to be flexibly allocated between competing demands for social interaction and sleep recovery.

Our results further indicate that only a brief, minimal period of social experience is sufficient to reshape subsequent sleep. A 24-hr period of social housing, even with only two flies per vial, was enough to increase sleep upon return to isolation, and this effect did not scale with group size over the interval tested. These observations raise the question of whether social experience operates as a binary signal (present/absent) or can scale with the amount or duration of social contact: does a minimal threshold of social exposure suffice to register social loss, or do shorter durations or sparser social configurations produce graded effects? Future experiments that further compress the duration of social housing or increase arena volume while keeping group size low could test whether the magnitude of the sleep response scales with social experience or instead reflects an all-or-none switch.

Several limitations should be considered. The social environment in sociSleep was intentionally simplified to enable long-term tracking of identified individuals, and the interaction metrics used here emphasized close proximity rather than the full repertoire of social behaviors. In addition, this study does not reveal the neural or molecular mechanisms through which recent social loss alters sleep and recovery. These questions are now tractable because social experience can be manipulated and measured without the confound introduced by assay-related state transitions.

## Methods and Materials

### *Drosophila* stock and maintenance

*w*CS were maintained under 12:12-hr light:dark (LD) cycles at 25 ℃ and ∼60% relative humidity. Male flies were collected within 24 hr of eclosion. In the experiments depicted in Figures 3 and 5, all flies had engaged in 3 to 5-day same-sex social interactions prior to being allocated to required housing conditions. In the experiments depicted in Figure 2A, flies were assigned to either single-housed (one fly per vial) or group-housed (six flies per vial) conditions after less than 24 hr of social interactions. In the experiments depicted in Figure 2D, to prevent effects from social-state transitions, singly housed flies were isolated since eclosion.

### sociSleep behavioral tracking system

Sleep and locomotion were monitored using a custom-built sociSleep system for long-term, identity-preserving tracking in solitary and social conditions. The system was housed in incubators under 12:12-hr light:dark cycles at 25 ℃ and ∼60% relative humidity. It consisted of a transparent 12-well suspension culture plate (Cat.-No. 665102, Greiner Bio-One) positioned above an infrared (IR) LED strip (940 nm, 12 V DC) with a diffuser sheet and recorded using an IR camera (Microsoft LifeCam Studio). Only the two central wells of the 12-well plate were used as sociSleep arenas, with one fly per well in solo trials or two flies per well in social trials. A physical air gap was maintained between the behavioral plate and the light source to reduce heating and stabilize arena temperature. Each arena was filled to ∼90% volume with agar-based food containing 5% sucrose, 1% tryptone, and 1% agar (all w/v), supplemented with 0.4% (v/v) propionic acid and 0.06% (v/v) phosphoric acid. Freshly prepared agar plates were left uncovered at room temperature for 3 hr before use.

Data acquisition and device control were performed on a Windows PC using Python scripts for solo-versus-social, solo-versus-solo, and social-versus-social experiments. The tracking pipeline outputs x-y coordinates and binary activity states (0, immobile; 1, moving) for each fly at 1-second resolution. SociSleep_tracker combines Kalman filter-based state prediction with nearest-neighbor assignment to preserve individual identities during tracking and incorporates arena-based prediction and detection constrains, merge-split event handling, and signal-loss recovery. The software includes a graphical interface to facilitate routine use. A demonstration of the workflow is provided (Movies S1 and S2).

### Sleep and locomotion quantification

Sleep was defined as immobility lasting more than 5 min, consistent with established criteria in *Drosophila* ^5^. Locomotor activity was quantified as second-to-second centroid displacement, with a movement threshold of 3 pixels/s used to distinguish locomotion from tracking noise and micro-movements. Sleep time and locomotor distance were calculated in 30-min bins using sociSleep_analyzer.py. Trajectories showing speed or co-movement were visualized and quantified using provided Python scripts. Co-movement was defined as periods of movement in which inter-fly distance was 30 pixels or less, used as a proxy for close-proximity interactions.

### Individual sleep measurement using ARC

Flies were transferred individually into the ARC system^16^ by mouth pipetting for automated sleep measurement. Recording tubes were arranged in alternating positions, with flies occupying every other tube (fly-empty-fly) to minimize potential visual or physical interactions between adjacent flies. Agar-based food, prepared as described for the sociSleep system, was loaded at the bottom of each tube to provide both nutrition and moisture.

### Housing geometry and encounter opportunity

To estimate whether social encounter opportunity differed between group-housing vials and sociSleep arenas, we calculated a density-normalized pairwise encounter index:

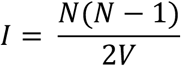

where *N* is fly number and *V* is accessible volume. This metric provides a coarse estimate of pairwise encounter opportunity based on group size and space and does not incorporate movement speed or interaction distance, which vary across housing conditions and circadian time. For the conditions used here, *N*_vial_ = 6, *N*_arena_ = 2, *V*_vial_ = 60*π* × 11^2^ mm^3^, and *V*_arena_ = 4*π* × 12^2^ mm^3^. Under this framework, pairwise interaction opportunities were comparable between group housing vials and sociSleep social arenas (I ≍ 6.5 × 10^-4^ mm^-3^ and 5.5 × 10^-4^ mm^-3^, respectively).

### Sleep deprivation

Sleep deprivation (Movie S10) was induced using intermittent 3D-mechanical stimulation^19^ from ZT12 to ZT24. Stimuli were delivered every 10 seconds for a 720° clockwise rotation followed by a vertical flick. Sleep recovery was quantified during the subsequent period (ZT0 – 24) using the sociSleep system.

### Data analysis

All statistical analyses were performed on paired measurements. Depending on the experiment, pairings were defined by matched conditions: solo vs. social, control vs. sleep-deprived, or solo vs. social sleep recovery. For social arenas containing two flies, values were averaged across individuals within each arena before statistical analysis. Sleep differences between conditions were evaluated using Wilcoxon signed-rank tests. Relationships between isolation duration and Δsleep (solo sleep – social sleep) were assessed using the Spearman’s rank correlation and robust linear regression. Wilcoxon signed-rank tests were used to determine if Δsleep was different from zero. Exact sample sizes and statistical tests are provided in the corresponding figure legends. All data analyses and visualizations were performed in Python (v3.12.0).

### Data availability

Movies S1-S10 and Table S1 are deposited in Zenodo (https://zenodo.org/records/21327147). The source code for the latest version of sociSleep is available at (https://github.com/HungryFly/sociSleep-for-fly-behavior). Programming language: Python. Operating system: Windows. Project name: sociSleep. GPL-3.0 open-source license.

## Acknowledgments

This work was funded by the NIH (R01DC020031, W.W.J.).

