## Supplementary Table 1 for "Recent social loss, not chronic isolation, reshapes sleep in *Drosophila*"

**Supplementary Table 1**: Manual validation of sociSleep performance in a social arena[1]

| Session ID (#) | Duration (min)[2] | Time stamp | Tracking error | Behavior[3] during errors | Classification[4] (sociSleep) | Fly | Sleep (min)[2] Manual | sociSleep |
|---|---|---|---|---|---|---|---|---|
| **1** | 10 | Day 1 ZT 6 | One identity swap | both moving | both moving | B | 5 | 0 |
| | | | | | | C | 0 | 5 |
| **2** | 13 | Day 2 ZT0 | / | | | B | 0 | 0 |
| | | | | | | C | 0 | 0 |
| **3** | 23 | Day 2 ZT 2 | One identity swap | both moving | both moving | B | 15 | 15 |
| | | | | | | C | 15 | 15 |
| **4** | 15 | Day 2 ZT 4 | / | | | B | 12 | 12 |
| | | | | | | C | 13 | 13 |
| **5** | 18 | Day 2 ZT 6 | / | | | B | 18 | 18 |
| | | | | | | C | 18 | 18 |
| **6** | 16 | Day 2 ZT 23 | / | | | B | 0 | 0 |
| | | | | | | C | 0 | 0 |
| **7** | 19 | Day 3 ZT 1 | Two signal losses (duration: 7s and 8s) | both immobile | both immobile | B | 19 | 19 |
| | | | | | | C | 18 | 18 |

[1]: The social arena contains two flies, designated FlyB and FlyC.

[2]: To simplify annotation, time is counted in minute-resolution here.

[3]: Manual annotation of fly behavior during error events (ground truth).

[4]: sociSleep-based automatic behavioral classification during error events.
